# scWeave: A deep learning model that bidirectionally translates between gene expression and chromatin structure at single-cell resolution

**DOI:** 10.64898/2026.07.13.738265

**Authors:** Ghulam Murtaza, Shengqi Hang, Xumeng Zhang, Shuhua Xu, Tangqi Fang, Doudou Yu, Anupama Jha, Ritambhara Singh, Sheng Wang, William Stafford Noble

## Abstract

Chromatin structure and gene expression are intimately linked, yet characterizing how the two covary has proven challenging, primarily because the two modalities are rarely measured in the same cells. Recently, single-cell co-assay protocols have enabled simultaneous profiling of both modalities within the same cells, but these experiments remain costly and technically challenging. To better characterize the relationship between 3D chromatin architecture and gene expression and to enable cross-modality inference from single-modality measurements, we developed a model called scWeave that bidirectionally translates between gene expression (scRNA-seq) and 3D chromatin architecture (scHi-C) at single-cell resolution. The scWeave model employs dual autoencoders to extract separate cell-level latent representations and learns to translate between these representations using dedicated translation modules. We evaluate scWeave on six publicly available co-assay datasets spanning mouse embryonic development, mouse cortex, mouse olfactory epithelium, and human bone marrow. On held-out mouse cells, scWeave outperforms a nearest-neighbor baseline and existing methods adapted to single-cell resolution, achieving a 57% improvement in median Spearman correlation when predicting gene expression from chromatin structure and an 18.8% improvement in median HiCRep similarity in the reverse direction relative to the next-best baseline. We further show that scWeave learns cross-modally aligned latent representations at single-cell resolution, enabling cells profiled in one modality to be matched to their counterparts in the other. Finally, scWeave generalizes to entirely held-out developmental timepoints in mouse olfactory epithelium and performs well on held-out human bone marrow cells despite limited human training data. By predicting the unmeasured chromatin architecture or transcriptional state from a single measured modality, scWeave offers a route to extend the benefits of costly co-assays to the many cell types, developmental stages, and species that are currently profiled with only one modality.

## 1 Introduction

One of the central questions in the study of 3D genome architecture is the extent to which the genome’s 3D conformation influences gene regulation. Chromatin is organized in a hierarchy of structures, including active and inactive compartments and topologically associating domains (TADs), which vary across cell types and developmental stages [1, 2]. It has been shown that these structures regulate gene expression by modulating the proximity between genes and their distal regulatory elements [3]. Hi-C [4] and RNA-seq [5] have emerged as the standard methods for profiling chromatin structure and gene expression, respectively, across bulk populations of cells. These two sequencing methods have opened the door to studying how chromatin structure and gene expression covary. Together, they have enabled researchers to investigate some of the most fundamental questions in genome biology, such as how a single genome gives rise to the many cell types that emerge during development and how dysregulation of this relationship drives disease [6–8]. However, characterizing how these two modalities relate and covary has proven challenging, in part because bulk data obscures mechanisms that operate at the single-cell or cell-type level.

Recently, a new class of experimental methods has emerged that enables the investigation of the relationship between gene expression and 3D genome architecture at single-cell resolution. Single-cell co-assays such as HiRES [2], GAGE-seq [9], LiMCA [10], CHARM [11], Paired Hi-C [12], and dscHi-C-multiome [13] produce, for each cell in a given dataset, a vector of gene expression values and a matrix of DNA–DNA contacts. By capturing both modalities within the same cell, these assays make it possible to directly link a cell’s genome organization to its transcriptional state. The resulting data reveals, for example, how the genome refolds as gene expression changes during differentiation and how structure, expression, and the relationship between them differ across cell types [2, 10]. However, such co-assays are costly and technically challenging [14], leaving many cellular and developmental states without paired measurements. Single-modality scRNA-seq and scHi-C experiments, by contrast, are far more abundant, but because they profile separate populations of cells, there is no direct way to know which cell in one experiment corresponds to which cell in the other or to recover the missing modality for a cell measured in only one.

One solution to this dearth of co-assay data is to train a machine learning model to predict the missing modality from a single measured modality. This approach has proved successful for other types of single-cell data. For example, existing cross-modality translation models can predict gene expression from chromatin accessibility (BABEL [15], Polarbear [16], and scButterfly [17]) and infer cell-surface protein abundance from gene expression (TotalVI [18] and sciPENN [19]). Beyond imputing missing modalities, these models learn joint representation spaces that can align cells across separately profiled, unpaired datasets. As a result, these methods have enabled atlas-scale integration of single-modality datasets and made it possible to jointly analyze cellular states across multiple modalities [20].

Accordingly, several recent methods aim to link scHi-C and scRNA-seq data (Table 1). scGrapHiC [21] predicts scHi-C from scRNA-seq using a deep neural network, while scHiGex [22] performs the reverse task, predicting scRNA-seq from scHi-C while additionally drawing on DNA sequence features. Because single-cell data is extremely sparse, both methods operate at the cell-type pseudobulk level, requiring cells to be grouped by cell type before training and inference. More recently, SEE [23] predicts scHi-C from scRNA-seq at single-cell resolution. However, SEE predicts contacts only within a short span around each gene and is consequently unable to predict the vast majority of chromatin contacts. Additionally, SEE requires cell-type labels during training to decide which scRNA-seq and scHi-C cells can be paired together. In practice, cell-type annotations are often noisy and inconsistent across experiments [24], and obtaining high-quality annotations is challenging when only scHi-C measurements are available.

**Table 1:**
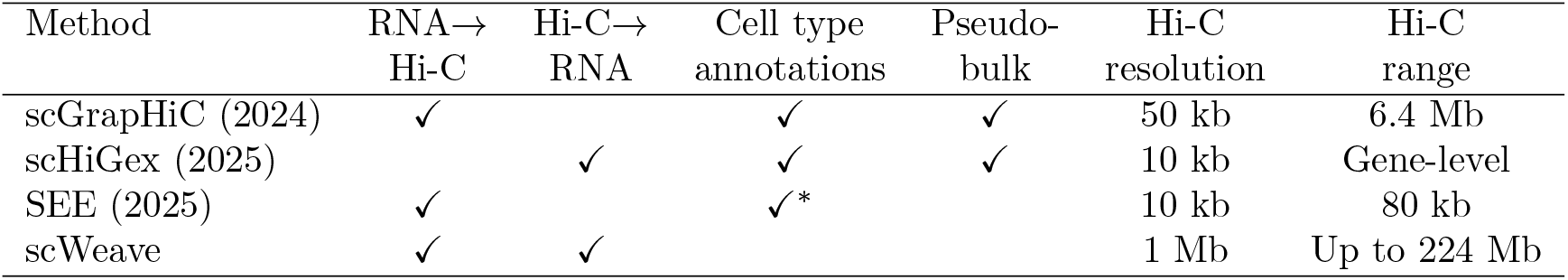
Comparison of approaches for translating between scRNA-seq and scHi-C. ^∗^SEE requires cell type annotation only for construction of the training set.

In this work, we propose to leverage co-assay data to train a machine learning model, scWeave, that bidirectionally translates between single-cell Hi-C (scHi-C) and single-cell RNA-seq (scRNA-seq) profiles. We designed the model around two goals. First, the model operates at single-cell rather than pseudobulk resolution, removing the need for cell-type annotations while preserving the cell-to-cell variation that pseudobulk aggregation discards. Second, scWeave predicts chromatin contacts across entire chromosomes rather than only over short genomic ranges, allowing the model to capture compartment- and chromosome-scale organization in addition to regional structure. The key challenge in meeting these goals is that single-cell data, especially scHi-C, is extremely sparse and high-dimensional, making direct prediction between raw modalities unreliable. Our model addresses this challenge in two ways. First, it encodes gene expression and chromatin structure into separate cell-level latent representations and translates between them using dedicated modules, so that cross-modality prediction occurs in a compact representation space rather than directly between raw feature spaces. Second, to obtain reliable structural representations from sparse single-cell contact matrices, scWeave builds on HiCFoundation [25], a foundation model pretrained on bulk Hi-C data.

We train and evaluate scWeave on six publicly available scRNA-seq and Hi-C co-assay datasets spanning mouse embryonic development, mouse cortex, mouse olfactory epithelium, and human bone marrow. On randomly held-out cells from the mouse embryonic development and brain cortex datasets, scWeave improves the median Spearman correlation for predicting gene expression from chromatin structure by 57% and the median HiCRep score [26], a distance-stratified similarity measure, for predicting chromatin structure from gene expression by 18.8%, relative to the next-best baseline. When evaluated on two entirely held-out developmental timepoints from mouse olfactory epithelium, scWeave improves the median Spearman correlation for gene expression prediction by 38% and the median HiCRep score for pseudobulk chromatin structure prediction by 27%, relative to the next-best baseline. Finally, when trained on human bone marrow cells, scWeave improves the median Spearman correlation for single-cell gene expression prediction by 98% and the median HiCRep score for pseudobulk chromatin structure prediction by 19%, relative to the next-best baseline. Our findings collectively suggest that scWeave can be applied across multiple species and tissue contexts to enable robust cross-modality translation. Beyond prediction, scWeave learns cross-modally aligned representations, placing each cell near its paired representation in the other modality. Cells profiled separately in each modality can then be matched based on these representations, and orthogonal epigenetic assays that the model does not use during training support the conclusion that these matches reflect biological similarity. Matching cells in this fashion can bring together large datasets that measure only gene expression or only chromatin structure, opening the door to joint analysis of transcriptional state and genome organization across cell types, developmental stages, and species for which only one modality has been profiled.

## 2 Results

### 2.1 scWeave accurately translates between gene expression and chromatin structure at single-cell resolution

The scWeave model consists of four main components: a gene expression autoencoder, a chromatin structure autoencoder, and two translation modules (RNA→Hi-C and Hi-C→RNA) that enable translation between gene expression and chromatin structure (Figure 1A). The modality-specific autoencoders encode each data type into dedicated cell-level latent representations. For scHi-C data, scWeave leverages the encoder from HiCFoundation, a chromatin structure foundation model trained on data from 494 bulk Hi-C experiments. We hypothesize that this pretrained encoder may facilitate stable and informative representation learning for chromatin structure. Furthermore, by encoding gene expression and chromatin structure independently and translating between their cell-level representations, scWeave avoids the difficult task of directly mapping between extremely high-dimensional raw feature spaces. This design choice simplifies the translation objective and allows the model to capture relationships between transcriptional state and genome organization.

**Figure 1:**
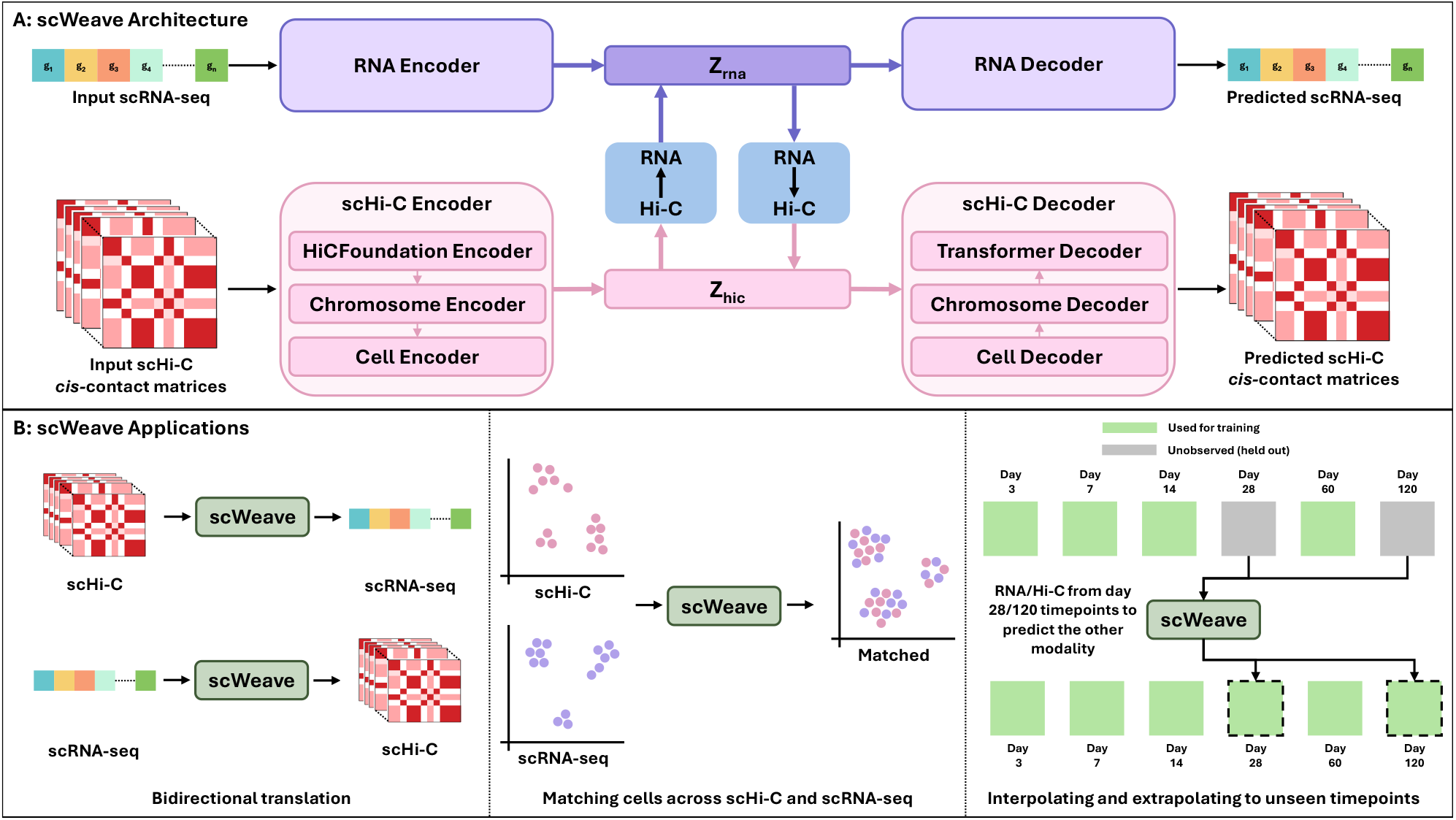
(A) **scWeave architecture.** scWeave comprises two modality-specific autoencoders together with two translation modules. The gene expression autoencoder encodes an input scRNA-seq profile (*g*_1_, …, *g*_*n*_) into a cell-level latent representation *Z*_*rna*_ and reconstructs the profile from it. The chromatin structure autoencoder encodes input scHi-C *cis*-contact matrices into a cell-level latent representation *Z*_*hic*_ and reconstructs them. The scHi-C encoder leverages a pretrained, frozen HiCFoundation encoder followed by chromosome- and cell-level encoders, and the decoder mirrors this architecture. The two translation modules (RNA→Hi-C and Hi-C→RNA) learn to map between *Z*_*rna*_ and *Z*_*hic*_, enabling bidirectional translation between the two modalities. (B) **scWeave applications**. (Left) Bidirectional translation: given a cell measured in one modality, scWeave predicts the other modality. (Middle) Cross-modality matching: cells profiled separately using scHi-C and scRNA-seq are matched by translating each cell into the target modality’s latent space and pairing it with its nearest neighbor in that space. (Right) Temporal interpolation and extrapolation: trained on a subset of measured developmental timepoints (green), scWeave can predict scRNA-seq from scHi-C and vice versa at held-out timepoints (gray), including an interpolated timepoint (day 28) between measured stages and an extrapolated timepoint (day 120) beyond them.

To validate scWeave’s ability to translate between modalities at single-cell resolution, we trained the model on co-assay datasets containing paired scRNA-seq and scHi-C measurements. These paired measurements enabled us to directly compare scWeave’s predictions with the corresponding observed modality in held-out cells. This experiment used mouse data from three studies, including HiRES data from 7,000 cells in mouse embryos (stages E7.0 through E15; HiRES-embryo) and 400 cells from the mouse cortex (HiRES-brain) [2], GAGE-seq data from 3,105 mouse cortex cells (GAGE-seq-brain) [9], and CHARM data from 4,265 mouse cortex cells (CHARM-brain) [11]. For each dataset, we randomly partitioned cells into training (80%), validation (10%), and test (10%). To our knowledge, no existing method can translate between gene expression and chromatin structure at single-cell resolution and generate chromosome-wide *cis*-contact predictions. Therefore, we compared scWeave against three baseline methods: a nearest-neighbor (NN) baseline that finds the closest-matching cell in the training dataset and uses its corresponding paired modality as the output, and two existing methods, scGrapHiC and scHiGex, which we adapted to operate at single-cell resolution. We retrained scGrapHiC and scHiGex on the same training datasets to ensure a fair comparison.

We first evaluated scWeave’s ability to predict gene expression from chromatin structure at single-cell resolution. We partitioned the test set by dataset and, for each of the three prediction methods (scWeave, NN, and scHiGex), compared the distribution of per-cell Spearman correlation coefficients (SCCs) between predicted and observed gene expression (Figure 2A). For this analysis, scGrapHiC is excluded because it only translates from gene expression to chromatin structure and not vice versa. Across all four datasets, which span three measurement protocols and two distinct tissue contexts, scWeave consistently achieves significantly higher scores than both baselines (*p* < 0.001, Wilcoxon signed-rank test). In particular, scWeave achieves a median score of 0.53, compared with 0.34 for NN and 0.07 for scHiGex, corresponding to a 57% relative improvement over the next-best baseline. The adapted scHiGex model performs poorly at single-cell resolution, with a median SCC of 0.07, which may reflect the difficulty of learning informative representations from extremely sparse scHi-C contact matrices. scWeave sidesteps this problem by building on HiCFoundation, whose pretrained chromatin structure representations provide greater robustness to sparse scHi-C contact matrices and enable accurate gene expression prediction.

**Figure 2:**
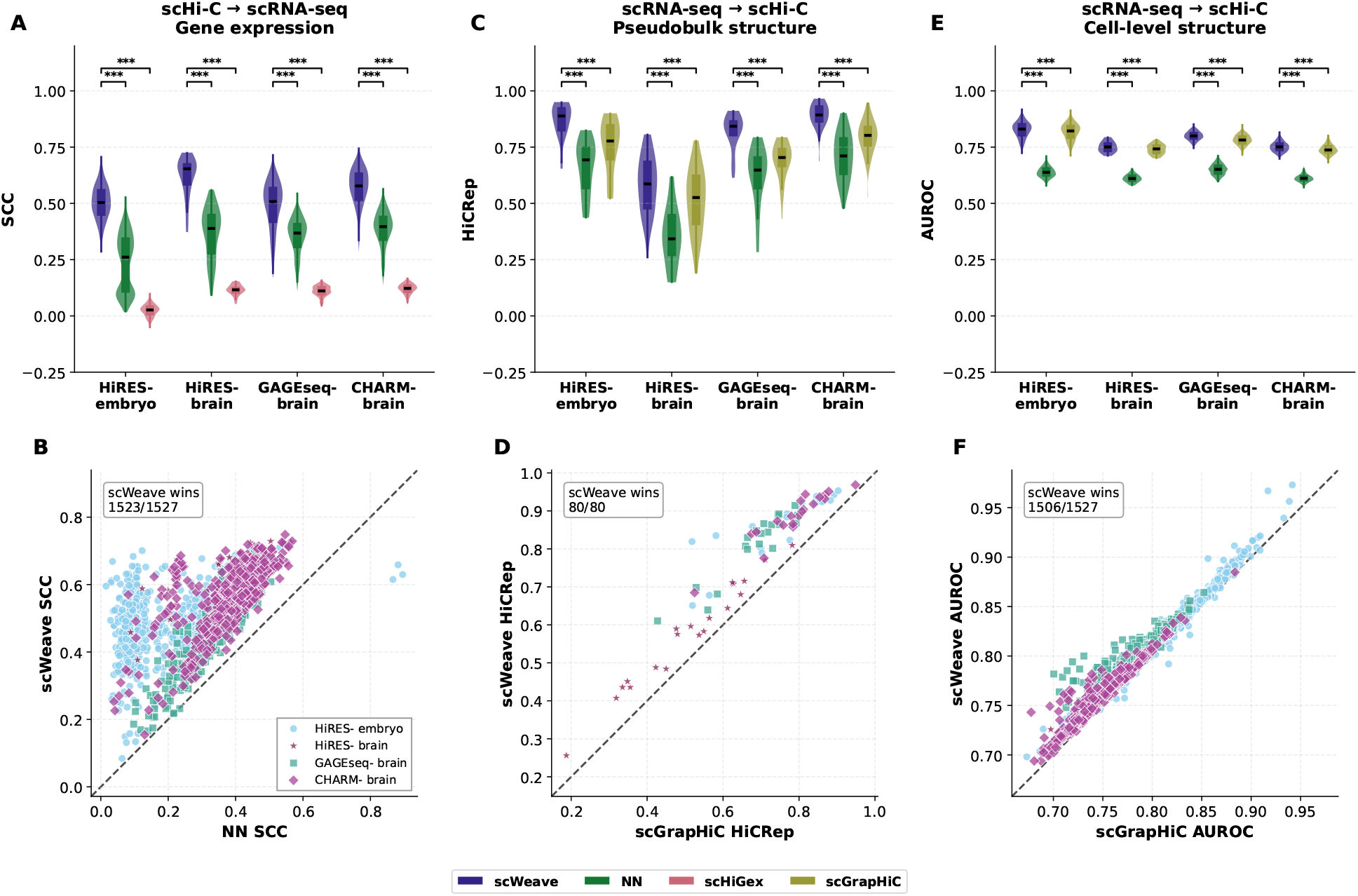
scWeave’s performance on bidirectional translation between gene expression and chromatin structure at single-cell resolution. (A) Violin plots showing the distribution of per-cell Spearman correlation coefficient (SCC) scores achieved on the Hi-C→RNA prediction task by each method (scWeave, NN, and scHiGex) in each dataset. Horizontal lines indicate the medians, and boxes indicate the interquartile ranges. Asterisks denote statistical significance, assessed using the one-sided Wilcoxon signed-rank test. Significance levels: * * * *p* < 0.001, ** *p* < 0.01, * *p* < 0.05. (B) Scatter plot in which each dot corresponds to a single cell, with the axes giving the SCC achieved on the Hi-C→RNA prediction task by the next-best baseline, NN (x-axis), and scWeave (y-axis). Points above the diagonal correspond to cells for which scWeave is more accurate. (C) Similar to panel A, but for the RNA→Hi-C prediction task evaluated at the pseudobulk level, using HiCRep as the performance measure and comparing scWeave, NN, and scGrapHiC. Because evaluation is performed at the pseudobulk level, each distribution is over chromosomes rather than single cells. (D) Similar to panel B, but for the RNA→Hi-C task evaluated at the pseudobulk level: each dot corresponds to a single chromosome in a single dataset, with the axes giving HiCRep for the next-best baseline, scGrapHiC (x-axis), and scWeave (y-axis). (E) Similar to panel A, but for single-cell RNA→Hi-C prediction evaluated as a binary classification task, using the area under the ROC curve (AUROC) as the performance measure and comparing scWeave, NN, and scGrapHiC. Each distribution is over cells. (F) Similar to panel B, but for single-cell RNA→Hi-C prediction: each dot corresponds to a single cell, with the axes giving AUROC for the next-best baseline, scGrapHiC (x-axis), and scWeave (y-axis).

To highlight performance across individual cells, we show a scatter plot (Figure 2B) comparing the per-cell Spearman correlation coefficients (SCCs) obtained by scWeave (y-axis) and the next-best baseline, NN (x-axis); points above the diagonal denote cells for which scWeave is more accurate. In this analysis, scWeave outperforms NN in 1,523 of 1,527 test-set cells, highlighting scWeave’s robust predictive accuracy across the entire test set. Of the four cells for which scWeave is less accurate than NN, three correspond to cases in which NN performs anomalously well. These three cells have extremely low scHi-C coverage, and we found that NN’s similarity-based search exploits this low coverage by retrieving training cells that share similar patterns of missingness rather than underlying biological structure.

Next, we evaluated the converse task of predicting chromatin structure from gene expression. We first carried out this evaluation at the pseudobulk level. To do so, we constructed pseudobulk Hi-C profiles by aggregating the test-set predictions within each dataset, yielding one predicted *cis*-contact matrix per chromosome, and separately constructed the corresponding pseudobulk profiles from the ground-truth data. We then compared the predicted and ground-truth *cis*-contact matrices using HiCRep. To compare the methods, we plotted the distributions of per-chromosome HiCRep scores as violin plots, stratified by dataset, for scWeave and the two baselines, NN and scGrapHiC (Figure 2C). For this analysis, scHiGex was excluded because it only predicts gene expression from chromatin structure. Across all four datasets, scWeave achieved the highest HiCRep scores, with its per-chromosome distributions shifted above those of both baselines, and these improvements were statistically significant in every dataset (*p* < 0.001, Wilcoxon signed-rank test). The advantage held for all chromosomes. Comparing the per-chromosome HiCRep scores of scWeave with those of the next-best baseline, scGrapHiC (Figure 2D), we observed that scWeave was more accurate on all 80 chromosomes. This observation shows that the improvement was consistent across individual chromosomes rather than reflecting only an aggregate trend.

In addition to the pseudobulk comparisons, we also evaluated chromatin structure prediction at single-cell resolution. Because single-cell Hi-C matrices are extremely sparse, comparing exact contact counts is not informative. Therefore, we reframed single-cell chromatin structure prediction as a binary classification task. For each cell, we ranked the predicted contacts by their scores and constructed a receiver operating characteristic (ROC) curve against the binarized ground truth, summarizing performance using the area under the curve (AUROC). We plotted the distributions of per-cell AUROC values as violin plots, stratified by dataset, for scWeave and the two baselines, NN and scGrapHiC (Figure 2E). Across all four datasets, scWeave consistently achieved the highest AUROC, with its per-cell distributions shifted above those of both baselines, and these improvements were statistically significant in every dataset (*p* < 0.001, Wilcoxon signed-rank test). The advantage also held in cell-by-cell comparisons. A paired scatter plot comparing scWeave with the next-best baseline, scGrapHiC (Figure 2F), shows that scWeave was more accurate in 1,506 of 1,527 test cells.

Together, these results show that scWeave accurately translates between gene expression and chromatin structure in both directions at single-cell resolution, consistently outperforming a nearest-neighbor baseline as well as retrained versions of the specialized models scHiGex and scGrapHiC across diverse assays and tissue contexts.

### 2.2 Orthogonal epigenetic assays confirm that scWeave learns biologically meaningful cross-modally aligned representations

Beyond predicting the held-out modality, we investigated how scWeave connects the two modalities through its learned representations. Specifically, we investigated whether each translation module maps a cell into the target modality’s latent space near its corresponding representation. For each cell in the test set, we extracted the translated latent representation (for example, the chromatin latent representation predicted from gene expression) and the ground-truth latent representation obtained by directly encoding the observed target modality. Visualization of the resulting embeddings using UMAP (Figure 3A) suggested that the two sets of points were closely intermingled. To quantify the quality of the alignment, we computed, for each cell, the fraction of samples closer than the true match (FOSCTTM) [27], where 0 indicates that a cell’s translated latent representation is closest to its own ground-truth counterpart and 0.5 indicates random alignment. To compare datasets of substantially different sizes, we sorted the per-cell FOSCTTM values in ascending order and plotted them against cell percentile (Figure 3B,C), showing that the vast majority of cells had low FOSCTTM values. Specifically, across the four datasets and both translation directions, the median FOSCTTM ranged from 0.004 to 0.021, indicating that scWeave mapped each cell into the other modality’s latent space near the location of its true partner.

**Figure 3:**
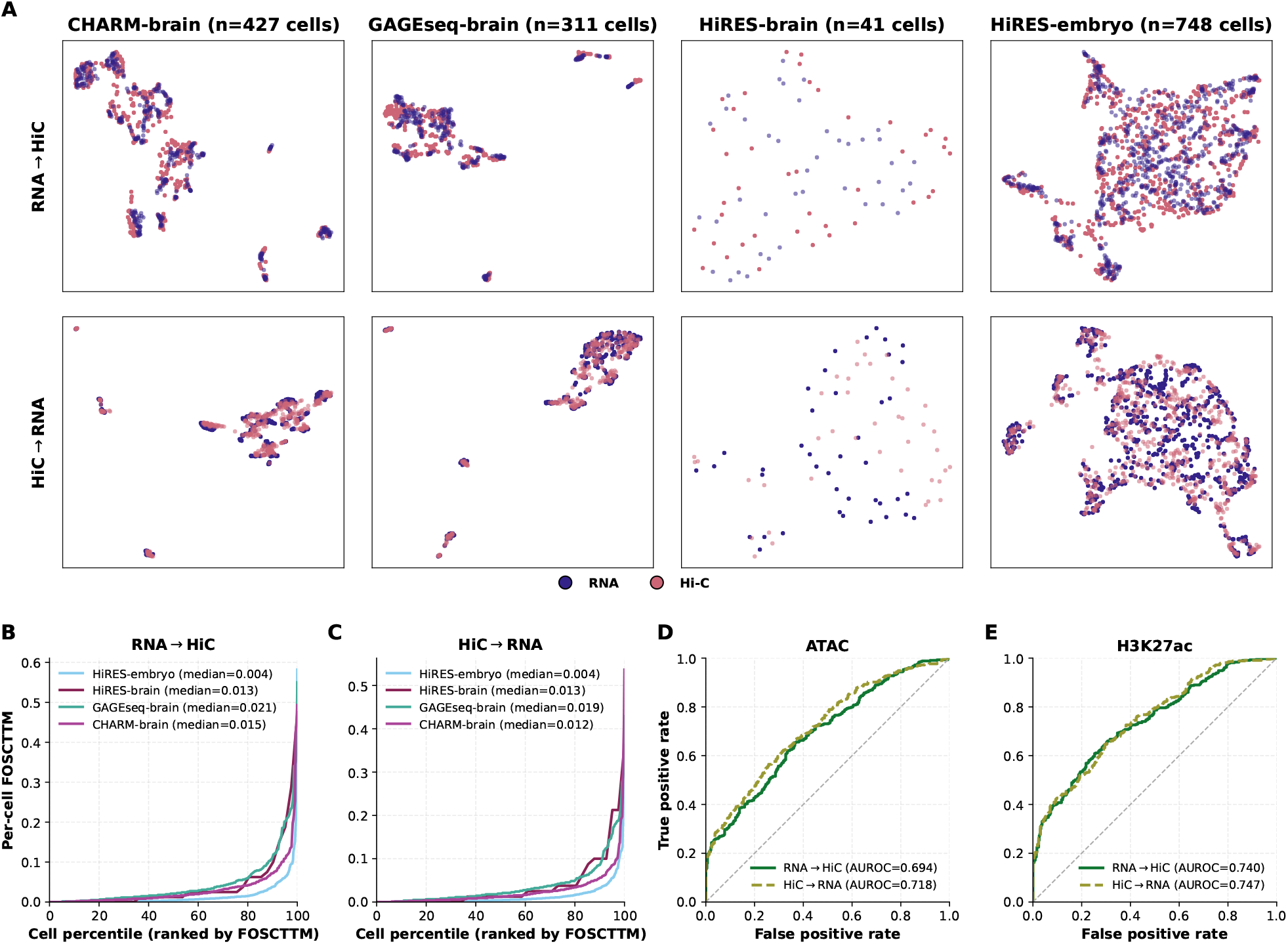
scWeave learns cross-modally aligned latent representations, as validated by orthogonal assays. (A) Joint UMAP embeddings of scWeave’s cell-level latent representations for each of the four held-out test datasets (columns). In each panel, the translated latent representations (indigo) are overlaid on the ground-truth encoded latent representations of the target modality (rose), with both jointly embedded using UMAP. The top row shows the RNA→Hi-C direction (predicted chromatin latent representations versus encoded chromatin latent representations), and the bottom row shows the Hi-C→RNA direction. (B, C) Per-cell fraction of samples closer than the true match (FOSCTTM) for the RNA→Hi-C (B) and Hi-C→RNA (C) directions, where 0 indicates that a cell’s true counterpart is its closest match and 0.5 indicates random alignment. For each dataset, the per-cell FOSCTTM values are sorted in ascending order and plotted against cell percentile (rank divided by the number of cells), allowing datasets of different sizes to be directly compared; the legend reports the median FOSCTTM for each dataset. (D, E) Orthogonal validation that the learned representations preserve single-cell biological similarity, shown for the CHARM-brain dataset using two assays not used during training: scATAC-seq (D) and H3K27ac (E). We use these assays as independent readouts of biological similarity. We quantify the similarity between cells using the Pearson correlation coefficient (PCC) between their scATAC-seq or H3K27ac profiles. We then frame the validation as a binary classification task scored using this PCC, where a positive example is a pair consisting of a query cell and its top-1 nearest neighbor in the target latent space, and a negative example is a randomly drawn cell pair. Each panel shows one ROC curve per translation direction (green solid, RNA→Hi-C; olive dashed, Hi-C→RNA) together with its AUROC; curves above the diagonal indicate that scWeave’s matched pairs are more similar in the orthogonal assays than randomly selected pairs, showing that latent-space matching retrieves epigenetically similar cells.

To further investigate the quality of the matches produced by scWeave, we leveraged the four-way co-assayed CHARM-brain data, which additionally profiles scATAC-seq and H3K27ac in the same cells. These two assays provide orthogonal readouts that were not used during model training. We therefore used the scATAC-seq and H3K27ac profiles to determine whether cells matched in scWeave’s latent space were biologically similar. We quantified this biological similarity by computing the Pearson correlation coefficient between cells using profiles from the same assay (scATAC-seq or H3K27ac). For each cell, scWeave produces a match by translating the cell into the other modality’s latent space and retrieving the cell whose ground-truth latent representation is nearest to the translated latent representation. For the RNA→Hi-C direction, for example, we encode the cell’s gene expression, predict its chromatin-structure latent representation, and retrieve the cell whose directly encoded Hi-C latent representation is nearest. We then framed the validation as a binary classification task. The positive examples are the matched pairs, each formed by a cell and the cell retrieved by scWeave as its match; the negative examples are an equal number of randomly drawn cell pairs. If scWeave’s matches are biologically meaningful, the matched pairs should have higher orthogonal-assay PCC values than the random pairs. For each assay and translation direction, we summarized this separation using an ROC curve and its area under the curve (AUROC), which represents the probability that a matched pair has a higher PCC than a random pair (Figure 3D,E). Across both assays and translation directions, the ROC curves lie well above the diagonal, with AUROC values ranging from 0.69 to 0.75. This separation is consistent across both assays and both directions, indicating that scWeave’s matches are biologically meaningful regardless of which modality is predicted or which epigenetic readout is used to assess similarity.

Together, these analyses show that scWeave learns cross-modally aligned latent representations at single-cell resolution and that their neighborhood structure reflects biological similarity, as supported by orthogonal measurements.

### 2.3 scWeave can accurately interpolate between and extrapolate to unseen timepoints in olfactory tissue growth

A common challenge in single-cell co-assay studies of developmental stages is that measurements are collected at only a limited number of timepoints because of cost, experimental throughput, and tissue availability. Consequently, many developmental stages remain unmeasured or are profiled using relatively inexpensive single-modality assays. Thus, there is a need for methods that infer missing modalities from observed data at these partially measured timepoints. Having demonstrated that scWeave can accurately perform bidirectional translation between gene expression and chromatin structure in held-out cells, we next tested whether these capabilities generalize across the transition from late developmental stages to young adulthood. For this experiment, we trained scWeave using the four previously described mouse co-assay datasets together with cells from four of the six LiMCA timepoints. To simulate partially observed timepoints, we used cells from four of the six available LiMCA timepoints (days 3, 7, 14, and 60) for training and validation, reserving measurements from postnatal days 28 and 120 for testing. We retained the previously described training, validation, and test splits for the HiRES-embryo, HiRES-brain, GAGE-seq-brain, and CHARM-brain datasets. The three cortex datasets were collected from 8–11-week-old mice. We extended the training set to include 90% of cells from the four selected LiMCA timepoints, adding the remaining 10% of cells to the validation set.

We first evaluated how well scWeave predicted gene expression from chromatin structure at single-cell resolution at timepoints that were not observed during training. Comparing the distributions of per-cell Spearman correlation coefficients (SCCs) between ground-truth gene expression and predictions from scWeave and the two baselines (NN and scHiGex), stratified by held-out timepoint (Figure 4A), we found that scWeave achieved the highest SCC at both timepoints. At day 28 (70 cells), scWeave attained a median SCC of 0.43, compared with 0.35 for NN and −0.04 for scHiGex; at day 120 (35 cells), it attained a median SCC of 0.52, compared with 0.34 for NN and −0.05 for scHiGex. At both timepoints, the improvement over each baseline was statistically significant (*p* < 0.001, Wilcoxon signed-rank test). To highlight performance across individual cells, we additionally compared the per-cell SCCs achieved by scWeave and the next-best baseline, NN, using a paired scatter plot (Figure 4B). We found that scWeave was more accurate in 69 of 70 cells at day 28 and 33 of 35 cells at day 120 (102 of 105 held-out cells in total), demonstrating robust predictive accuracy at the level of individual cells across both held-out timepoints. As observed previously, the adapted scHiGex model performed poorly at single-cell resolution, showing a near-zero or slightly negative median SCC (−0.04 at day 28 and −0.05 at day 120) relative to the ground-truth gene expression.

**Figure 4:**
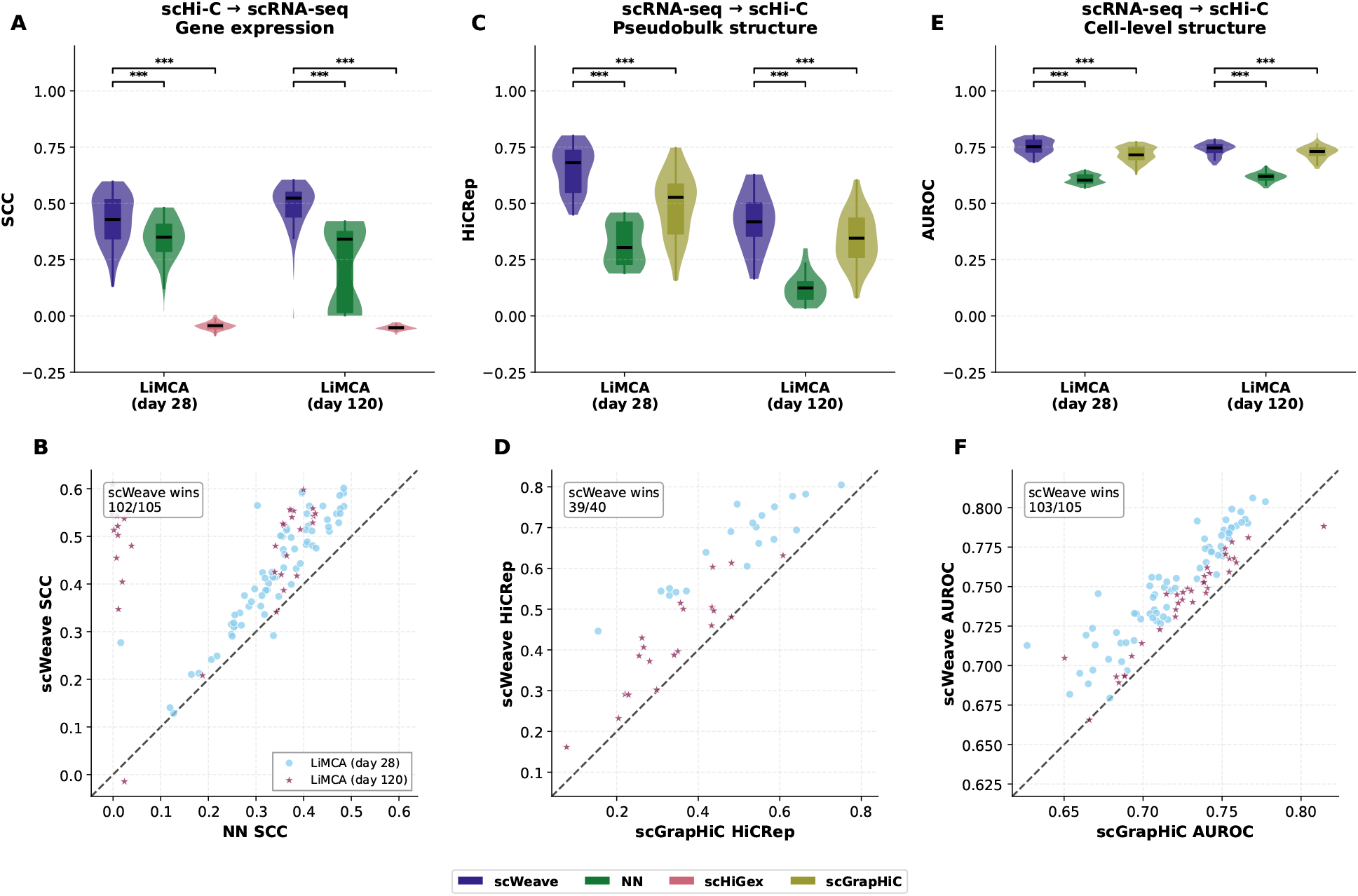
scWeave’s performance on held-out timepoints (day 28 and day 120) in LiMCA mouse olfactory tissue. (A) Violin plots showing the distributions of per-cell Spearman correlation coefficient (SCC) scores achieved on the Hi-C→RNA prediction task by each method (scWeave, NN, and scHiGex) for cells from day 28 and day 120 in the LiMCA mouse dataset. Horizontal lines indicate the medians, and boxes indicate the interquartile ranges. Asterisks denote statistical significance, assessed using the one-sided Wilcoxon signed-rank test. Significance levels: *** *p* < 0.001, ** *p* < 0.01, * *p* < 0.05. (B) Scatter plot in which each dot corresponds to a single cell, with the axes giving the SCC achieved on the Hi-C→RNA prediction task by the next-best baseline, NN (x-axis), and scWeave (y-axis). Points above the diagonal correspond to cells for which scWeave is more accurate. (C) Similar to panel A, but for the RNA→Hi-C prediction task evaluated at the pseudobulk level, using HiCRep as the performance measure and comparing scWeave, NN, and scGrapHiC. Because evaluation is performed at the pseudobulk level, each distribution is over chromosomes rather than single cells. (D) Similar to panel B, but for the RNA→Hi-C task evaluated at the pseudobulk level: each dot corresponds to a single chromosome at one timepoint, with the axes giving HiCRep for the next-best baseline, scGrapHiC (x-axis), and scWeave (y-axis). (E) Similar to panel A, but for single-cell RNA→Hi-C prediction evaluated as a binary classification task, using the area under the ROC curve (AUROC) as the performance measure and comparing scWeave, NN, and scGrapHiC. Each distribution is over cells. (F) Similar to panel B, but for single-cell RNA→Hi-C prediction: each dot corresponds to a single cell, with the axes giving AUROC for the next-best baseline, scGrapHiC (x-axis), and scWeave (y-axis).

Next, we evaluated the converse direction, predicting chromatin structure from gene expression at the held-out timepoints, at both the pseudobulk and single-cell levels. At the pseudobulk level, we aggregated the test-set predictions at each timepoint into one predicted *cis*-contact matrix per chromosome and compared them with the ground-truth pseudobulk profiles using HiCRep (Figure 4C). scWeave achieved the highest median HiCRep at both timepoints (0.68 at day 28 and 0.42 at day 120, compared with 0.53/0.35 for scGrapHiC and 0.28/0.12 for NN; *p* < 0.001, Wilcoxon signed-rank test) and was more accurate than the next-best baseline, scGrapHiC, on 39 of 40 chromosomes (Figure 4D). At the single-cell level, when prediction was framed as a binary classification task scored using AUROC (Figure 4E), scWeave again achieved the highest median AUROC at both timepoints (0.75 at both day 28 and day 120, compared with 0.72/0.73 for scGrapHiC and 0.60/0.62 for NN; *p* < 0.001) and outperformed scGrapHiC in 103 of 105 held-out cells (Figure 4F). Notably, scWeave’s pseudobulk HiCRep was lower at the extrapolated day-120 timepoint than at the interpolated day-28 timepoint (0.42 versus 0.68), a decline that we did not observe when predicting gene expression from chromatin structure (Figure 4A).

Collectively, our results show that scWeave can accurately interpolate chromatin structure and gene expression at intermediate, partially observed developmental timepoints and extrapolate from developmental stages to young adulthood.

### 2.4 scWeave outperforms existing baselines on bidirectional translation in a data-limited human bone marrow dataset

Most available co-assayed scRNA-seq and scHi-C data to date has been collected from mouse samples. To test whether scWeave could learn meaningful associations between chromatin structure and gene expression from the limited available human data, we trained scWeave and the existing baselines from scratch on co-assayed scRNA-seq and scHi-C measurements from human bone marrow profiled using the GAGE-seq protocol [9]. This dataset comprises only 1,097 cells, which we partitioned into 80% training, 10% validation, and 10% test sets, leaving only 877 cells for training. For comparison, the mouse model in Figure 2 was trained on 12,190 cells, approximately 14 times as much training data drawn from a substantially more diverse set of datasets. All models were trained entirely on human data without transferring any weights from the mouse models, thereby avoiding the need to map orthologs between species.

In this setting, scWeave achieved the highest performance in both translation directions. Specifically, for the Hi-C→RNA task, scWeave achieved a median per-cell SCC of 0.30, compared with 0.15 for NN and 0.01 for scHiGex (Figure 5A), and was more accurate than the next-best baseline, NN, in all 111 held-out cells (Figure 5B). For the converse RNA→Hi-C task, scWeave achieved a median pseudobulk HiCRep of 0.64 (compared with 0.54 for NN and 0.16 for scGrapHiC; Figure 5C) and a median cell-level AUROC of 0.82 (compared with 0.78 for scGrapHiC and 0.64 for NN; Figure 5E), outperforming the next-best baseline on 22 of 23 chromosomes and all 111 cells, respectively (Figure 5D and F). All improvements over both baselines were statistically significant (*p* < 0.001, Wilcoxon signed-rank test).

**Figure 5:**
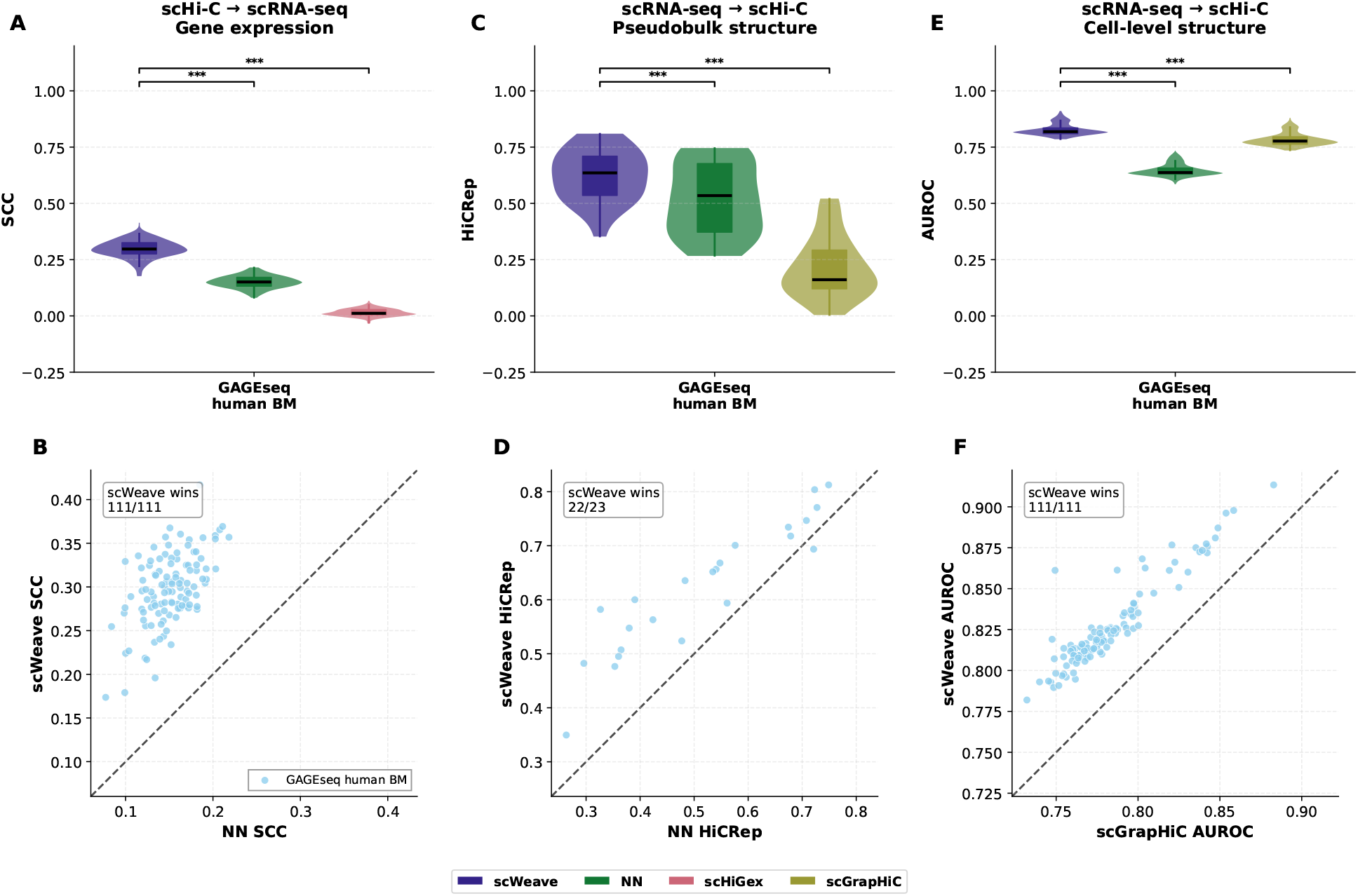
scWeave’s performance on held-out human bone marrow cells. (A) Violin plot showing the distribution of per-cell Spearman correlation coefficient (SCC) scores achieved on the Hi-C→RNA prediction task by scWeave, NN, and scHiGex (with methods distinguished by color) across all held-out human bone marrow cells. Horizontal lines indicate the medians, and boxes indicate the interquartile ranges. Asterisks denote statistical significance, assessed using the one-sided Wilcoxon signed-rank test. Significance levels: *** *p* < 0.001, ** *p* < 0.01, * *p* < 0.05. (B) Scatter plot in which each dot corresponds to a single cell, with the axes giving the SCC achieved on the Hi-C→RNA task by the next-best baseline, NN (x-axis), and scWeave (y-axis). Points above the diagonal correspond to cells for which scWeave is more accurate. (C) Similar to panel A, but for the RNA→Hi-C prediction task evaluated at the pseudobulk level, using HiCRep as the performance measure and comparing scWeave, NN, and scGrapHiC. Because evaluation is performed at the pseudobulk level, the distribution is over chromosomes rather than single cells. (D) Similar to panel B, but for the RNA→Hi-C task evaluated at the pseudobulk level: each dot corresponds to a single chromosome, with the axes giving HiCRep for the next-best baseline, NN (x-axis), and scWeave (y-axis). (E) Similar to panel A, but for single-cell RNA→Hi-C prediction evaluated as a binary classification task, using the area under the ROC curve (AUROC) as the performance measure and comparing scWeave, NN, and scGrapHiC. The distribution is over cells. (F) Similar to panel B, but for single-cell RNA→Hi-C prediction: each dot corresponds to a single cell, with the axes giving AUROC for the next-best baseline, scGrapHiC (x-axis), and scWeave (y-axis).

Despite consistently outperforming the baselines, scWeave’s absolute performance on human data was lower than on mouse data. Relative to its performance on mouse data generated using the same GAGE-seq protocol, scWeave’s median SCC for gene-expression prediction was 41% lower (0.30 versus 0.51), and its median pseudobulk HiCRep was 25% lower (0.64 versus 0.84). With only a single, relatively small human dataset, it is difficult to attribute this gap to any single cause. This reduction in predictive performance could be attributed to the limited amount of human training data, the intrinsic difficulty of predicting human chromatin structure, systematic technical differences between the mouse and human datasets, or some combination of these factors.

Even in this constrained data regime, scWeave achieved the highest scores across both translation directions and all three metrics compared with the baselines. This result suggests that scWeave’s dual-autoencoder design with dedicated modality-translation modules provides an effective approach for learning mappings between gene expression and chromatin structure, even when the available data is extremely limited.

## 3 Discussion

In this work, we introduced scWeave, a model that learns to translate bidirectionally between gene expression (scRNA-seq) and chromatin structure (scHi-C) at single-cell resolution. We validated scWeave across multiple complementary evaluation settings to assess both predictive accuracy and generalization. In randomly held-out cells from four independent mouse co-assay datasets, scWeave consistently outperformed a nearest-neighbor (NN) baseline and the existing pseudobulk methods scHiGex and scGrapHiC, which we repurposed and retrained for single-cell prediction. We further showed that scWeave learns cross-modally aligned representations at single-cell resolution and used orthogonal epigenetic assays to confirm that similarity within these representations reflects biological similarity. We also evaluated scWeave at entirely held-out developmental timepointsin the mouse olfactory epithelium and found that the model successfully interpolated between measured stages while retaining substantial predictive power when extrapolating to an entirely held-out developmental timepoint. Finally, although the model trained on the limited human co-assay data was less accurate than the mouse model, it still outperformed competing baselines on held-out human bone marrow cells. Collectively, these results show that scWeave accurately and robustly translates between transcriptional state and chromatin structure at single-cell resolution while demonstrating strong generalizability.

Although scWeave demonstrates strong bidirectional translation between gene expression and chromatin structure, several limitations remain. First, its reliance on HiCFoundation constrains scWeave to operate at 1 Mb resolution and to inherit architectural and chromatin-structure assumptions from a model trained on bulk Hi-C data, which may limit its ability to capture single-cell-specific structural features. Accordingly, fine-tuning HiCFoundation on a large compendium of scHi-C data could improve the model’s performance and allow it to operate at a finer resolution. Second, despite the robust embeddings provided by HiCFoundation, single-cell Hi-C data remains extremely sparse. In this work, we focused on co-assays that yield relatively deep scHi-C coverage and excluded assays such as dscHi-C-multiome and Paired Hi-C, which yield very low contact depths (median contact counts of ~45,000 and ~33,000 per cell, respectively). One approach to addressing this sparsity would be to incorporate DNA sequence features alongside sparse chromatin-structure measurements to further improve the robustness of the chromatin-structure representations. Third, the availability of co-assayed scRNA-seq and scHi-C datasets remains limited in both scale and diversity, severely restricting the training data available to scWeave. As a potential next step, we could pretrain the autoencoders using the much more widely available single-modality scHi-C and scRNA-seq data to further improve performance and generalization. Overall, scWeave provides a foundation for integrating transcriptional and structural modalities at single-cell resolution and establishes a framework for studying the coordinated relationships between gene expression and chromatin structure.

## 4 Methods

### 4.1 Datasets and preprocessing

To train and test scWeave, we gathered six publicly available scRNA-seq and scHi-C co-assay datasets from four studies (Table 2). The first study employed the HiRES protocol to generate scRNA-seq and scHi-C data for over 7000 cells from mouse embryos spanning developmental stages E7.0 to E15, along with approximately 400 cells from the mouse cortex [2]. The second study profiled 3105 cells from the mouse cortex using the GAGE-seq protocol [9]. The GAGE-seq study also profiled 1097 human bone marrow cells, which we used to train and evaluate scWeave on human data. The third study used the CHARM protocol to profile 4265 cells from the mouse cortex [11]. The last study used the LiMCA protocol to collect data from 411 cells in the mouse olfactory epithelium across six time points (postnatal days 3, 7, 14, 28, 60, and 120) [10]. For the temporal generalization experiment, we completely held out LiMCA days 28 and 120 to evaluate scWeave’s bidirectional translation performance in interpolation and extrapolation settings. Unless otherwise stated, each dataset is randomly partitioned into 80% training, 10% validation, and 10% test splits.

**Table 2:** Single-cell RNA/Hi-C co-assay datasets. The table lists the number of cells and the median numbers of Hi-C contacts and RNA-seq UMIs per cell. LiMCA UMIs are reported as N/A because its scRNA-seq protocol quantifies gene expression by read count rather than byunique molecular identifiers (UMIs).

| Assay | Species | Tissue | Cells | Cell types | Contacts | UMIs |
| --- | --- | --- | --- | --- | --- | --- |
| HiRES | mouse | developing embryo | 7469 | 21 | 306,194 | 9,865 |
| HiRES | mouse | brain cortex | 399 | 7 | 376,633 | 28,709 |
| GAGE-seq | mouse | brain cortex | 3105 | 29 | 393,445 | 20,160 |
| GAGE-seq | human | bone marrow | 1097 | 6 | 400,952 | 5,504 |
| CHARM | mouse | brain cortex | 4265 | 20 | 485,033 | 8,959 |
| LiMCA | mouse | olfactory epithelium | 411 | 5 | 652,291 | N/A |

For all datasets, we applied a common series of preprocessing steps. For the scRNA-seq data, we first normalized each cell’s counts to a total of 10,000, then applied a log1p transformation (natural logarithm of counts plus one) for variance stabilization. Additionally, for each species, we retained only genes expressed in at least 20% of the training cells, pooled across all datasets for that species. After applying this filter, we were left with 5656 mouse genes and 3037 human genes. For the scHi-C data, we applied the transformations required by HiCFoundation. First, for each chromosome, we aggregated its *cis*-contacts into a contact matrix at 1 Mb resolution. Second, we applied a log1p transformation. Third, we normalized the values by dividing each entry by the log-transformed maximum value, thereby ensuring that the matrix values are scaled between 0 and 1.

### 4.2 scWeave

The scWeave model consists of a gene expression autoencoder, a chromatin structure autoencoder, and two translation modules (RNA→Hi-C and Hi-C→RNA) that collectively support bidirectional translation between gene expression and chromatin structure.

#### 4.2.1 Gene expression autoencoder

The gene expression autoencoder consists of two modules: the encoder and the decoder. These modules jointly learn to encode log-normalized gene expression into a cell-level representation and then reconstruct the gene expression values from that representation. The encoder network *ϕ*_*rna*_ takes as input a log-normalized gene expression vector and projects it into a 2048-dimensional cell-level representation *Z*_*rna*_. The encoder network *ϕ*_*rna*_ comprises two fully connected layers with root mean square layer normalization (RMSNorm) [28] and a sigmoid linear unit (SiLU) activation [29] between them. The first layer maps the input gene expression vector *X*_*rna*_ to a hidden vector of dimension 4096, and the second layer maps this hidden vector to the latent vector *Z*_*rna*_ of dimension 2048. We then pass this latent vector *Z*_*rna*_ to a decoder network *θ*_*rna*_, which mirrors the encoder architecture in reverse, mapping *Z*_*rna*_ back to a reconstructed gene expression vector of the original input dimension.

#### 4.2.2 Chromatin structure autoencoder

Single-cell Hi-C measurements are substantially sparser and noisier than scRNA-seq. To mitigate this sparsity, we leverage an existing Hi-C foundation model, HiCFoundation, to obtain patch-level embeddings of chromosome-level *cis*-contact matrices for all autosomal chromosomes and the X chromosome at 1 Mb resolution. HiCFoundation expects a fixed input of size 224×224, which in our case encapsulates a single chromosome’s *cis*-contact matrix at 1 Mb resolution. Chromosomes shorter than 224 Mb are zero-padded to this size, while the few chromosomes longer than 224 Mb (human chromosomes 1 and 2) are cropped to their first 224 bins. HiCFoundation partitions each 224×224 input into non-overlapping 16×16 patches and produces a 1024-dimensional embedding for each patch, yielding a tensor of shape (*N*, 14, 14, 1024), where *N* is 20 for mouse and 23 for human. This tensor of patch-level representations is passed to a hierarchical autoencoder to first combine them into a single cell-level representation and from that cell-level representation decode *cis*-contact matrices for all chromosomes.

The encoder consists of two sub-modules, *ϕ*_*chrom*_ and *ϕ*_*cell*_. The *ϕ*_*chrom*_ module, shared across chromosomes, first projects each of a chromosome’s 196 patch embeddings from 1024 to 256 dimensions, then aggregates the resulting 196×256 patch-level embeddings into a single 1024-dimensional chromosome-level representation via two fully connected layers with a SiLU activation between them. Next, *ϕ*_*cell*_ further aggregates all 1024-dimensional chromosome-level representations by using a network with two fully connected layers with a SiLU activation between them to produce a single cell-level representation, *Z*_*hic*_, of dimension 2048. The decoder mirrors this hierarchy. The first decoder submodule, cell decoder *θ*_*cell*_, reconstructs chromosome-level embeddings from *Z*_*hic*_ by essentially performing an inverse operation of cell encoder *ϕ*_*cell*_. Next, reconstructed chromosome-level representations are passed to chromosome decoder *θ*_*chrom*_, which yields patch-level representations for each chromosome of size 196×256. Finally, scWeave passes the reconstructed patch-level representations of each chromosome to a shared lightweight vision transformer decoder, *θ*_*contact*_, that is composed of eight transformer blocks, each with eight attention heads. The transformer decoder maps each chromosome’s 196×256-dimensional embeddings into full 224×224 *cis*-contact matrices at 1 Mb resolution. During training, we kept the HiCFoundation encoder frozen and optimize the weights of all other components of the chromatin structure autoencoder.

#### 4.2.3 Bidirectional translation between gene expression and chromatin structure

To enable bidirectional translation between gene expression and chromatin structure we leverage two translation modules ζ_*rna*→*hic*_ and ζ_*hic*→*rna*_ that operate in the cell-level representation space. The ζ_*rna*→*hic*_ module is trained to predict *Z*_*hic*_ from *Z*_*rna*_, and the ζ_*hic*→*rna*_ module is trained to predict *Z*_*rna*_ from *Z*_*hic*_. Each translation module consists of two fully connected layers, each followed by RMS layer normalization, with a SiLU activation applied after the first layer, and a residual connection that adds the module’s input to its output. All layers operate at the cell-level representation dimension of 2048, so the hidden dimension matches the input for both RNA and Hi-C.

### 4.3 Loss function

Finally, to jointly train the two autoencoders along with the bidirectional translation modules, we minimize a combined loss

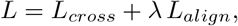

where *λ* = 0.1. The combined reconstruction and cross-modal translation loss *L*_*cross*_ is

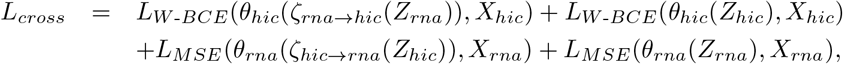

where *θ*_*hic*_ = *θ*_*contact*_ ◦ *θ*_*chrom*_ ◦ *θ*_*cell*_ denotes the full chromatin-structure decoder. *L*_*cross*_ ensures that scWeave learns to predict one modality from the other while also ensuring that both the gene expression and chromatin structure autoencoders preserve their ability to reconstruct their own modality. The mean squared error loss *L*_*MSE*_ optimizes reconstruction and accurate translation of gene expression, while *L*_*W* - *BCE*_ is a weighted binary cross-entropy loss that assigns a three times higher penalty on mispredicting non-zero values, to optimize reconstruction of scHi-C contact matrices. We chose binary cross-entropy loss for chromatin structure because single-cell Hi-C contact matrices are extremely sparse and are dominated by zeros, which consequently makes prediction of contact presence more important than precise regression of contact frequency. For the loss calculation, we binarized the normalized input scHi-C matrices by setting bins with no observed contacts to 0 and those with non-zero observed contacts to 1.

The latent alignment loss *L*_*align*_ encourages each translated representation to lie close to its corresponding representation in the target modality’s latent space. Let 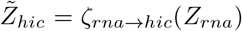 and 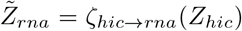 denote the translated latent representations. For a batch of *N* paired cells, we define the temperature-scaled cosine-similarity matrix as

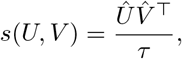

where *Û* and 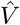 are the row-wise L2-normalized latent matrices and *τ* = 0.1 is the temperature parameter. Each entry *s*(*U, V*)_*ij*_ therefore represents the cosine similarity between cell *i* in *U* and cell *j* in *V*.

For a pair of latent matrices *U* and *V*, we define a symmetric contrastive matching loss that identifies each cell’s true paired counterpart among the cells in the batch. Let **y** = (1, …, *N*) denote the target indices, such that row *i* of the similarity matrix is matched to column *i*. The contrastive loss is

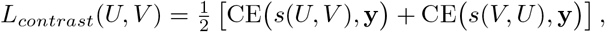

where CE denotes the cross-entropy loss. We apply this loss in both translation directions and average the resulting losses:

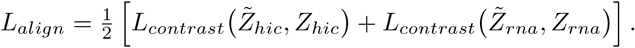

This objective increases the similarity between each cell’s translated latent representation and its corresponding ground-truth representation in the target modality while decreasing its similarity to the representations of other cells in the batch.

### 4.4 Baselines

To our knowledge,no existing method can predict chromosome-wide *cis*-contact matrices from gene expression, or gene expression from chromatin contact matrices, at single-cell resolution. We therefore compared scWeave against three baselines: a nearest-neighbor baseline and two existing methods that we repurposed to operate at single-cell resolution.

The nearest-neighbor (NN) baseline operates in a cross-modality setting. For a given input cell, it identifies the closest-matching cell in the training set based on the input modality and returns that training cell’s corresponding target modality as its prediction. Because both scRNA-seq and scHi-C data are extremely sparse and high-dimensional, we first embed gene expression and contact matrices into a 2048-dimensional PCA space (matching the dimensionality of scWeave’s cell-level latent representation) and then identify the nearest neighbor using cosine distance in that space.

In addition to the NN baseline, we repurposed two existing methods, scGrapHiC [21] and scHiGex [22], to operate at single-cell resolution. scGrapHiC takes single-cell gene expression as input and predicts the corresponding single-cell chromatin contact matrix, whereas scHiGex takes single-cell chromatin structure as input and predicts the corresponding single-cell gene expression. We retrained both methods on the same training datasets used for scWeave, using the hyperparameters reported in their original studies. We emphasize that both methods were originally designed for pseudobulk, cell-type-level prediction, so repurposing them for single-cell prediction falls outside their intended use case, and their performance here should not be taken as a reflection of how well they perform in their intended use case. Moreover, unlike scWeave, neither method can perform bidirectional translation: scGrapHiC only predicts chromatin structure from gene expression, and scHiGex only predicts gene expression from chromatin structure. Additionally, neither method can match cells across the gene expression and chromatin structure modalities.

### 4.5 Evaluation metrics

To assess how accurately scWeave translates between gene expression and chromatin structure, we compared model predictions to ground-truth measurements using a metric appropriate to each task.

#### Gene expression prediction

We evaluated gene-expression predictions at the single-cell level using the Spearman correlation coefficient (SCC) between the predicted and ground-truth gene expression vectors. We computed this correlation over all genes, including lowly expressed ones, to establish that scWeave accurately predicts cell-level gene expression.

#### Chromatin structure prediction

Because single-cell Hi-C data is extremely sparse, we evaluated chromatin structure predictions at both the single-cell and pseudobulk levels. At the single-cell level, we framed prediction as a binary classification task: we binarized the ground-truth contact matrices and, for each cell, constructed a receiver operating characteristic (ROC) curve of the predicted contacts against the binarized ground truth, summarizing it by the area under the curve (AUROC). We excluded very short-range contacts (the first three diagonals, ≤ 3 Mb) from this evaluation. At the pseudobulk level, we aggregated the predicted and ground-truth contact maps by averaging the per-cell contact matrices within each dataset and measured their similarity using HiCRep [26], a stratum-adjusted correlation that captures agreement in higher-order chromatin organization across genomic distances.

### 4.6 Implementation details

The entire pipeline, including scWeave, baselines, and evaluation metrics, is implemented in Python (version 3.12.13). scWeave is implemented with PyTorch Lightning (version 2.6.1) using the PyTorch (version 2.12.0) backend. scWeave uses HiCFoundation (version 1.0.0) with weights that are publicly available on the Hugging Face library. We trained scWeave for 25 epochs with a batch size of 8 and optimized the weights using the Adam optimizer with an initial learning rate of 1 × 10^−4^ and default values for the remaining optimizer parameters. For inference and testing we used the weights that achieved the minimum loss on the validation set. Training loss curves are provided in Supplementary Figure S1-S3.

## Code Availability

scWeave is open source under the Apache 2.0 License. The code for data processing, model training and model inference is publicly available on GitHub at (https://github.com/Noble-Lab/scWeave).

## Data availability

All single-cell RNA-seq and single-cell Hi-C data used in this manuscript is publicly available. The HiRES dataset is available at GEO GSE223917; the GAGE-seq dataset is available at GEO GSE238001; the CHARM dataset is available at GEO GSE303006 and the LiMCA dataset is available at GSE239969.

## Acknowledgements

This work was supported by NIH award R01 HG013321 and an NVIDIA compute grant.

## Author contributions

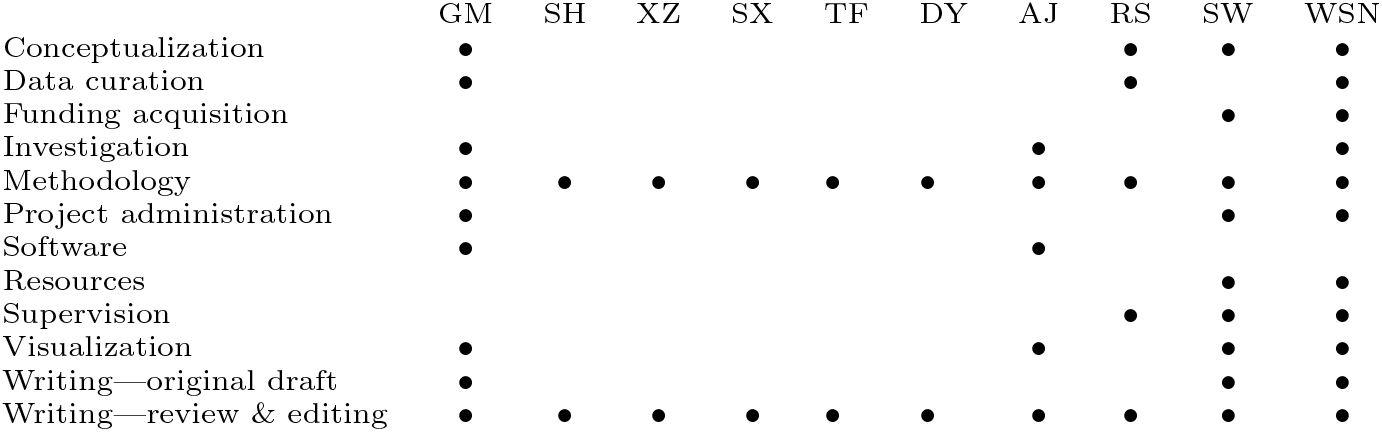

