## supplementary figure 1 for "scWeave: A deep learning model that bidirectionally translates between gene expression and chromatin structure at single-cell resolution"

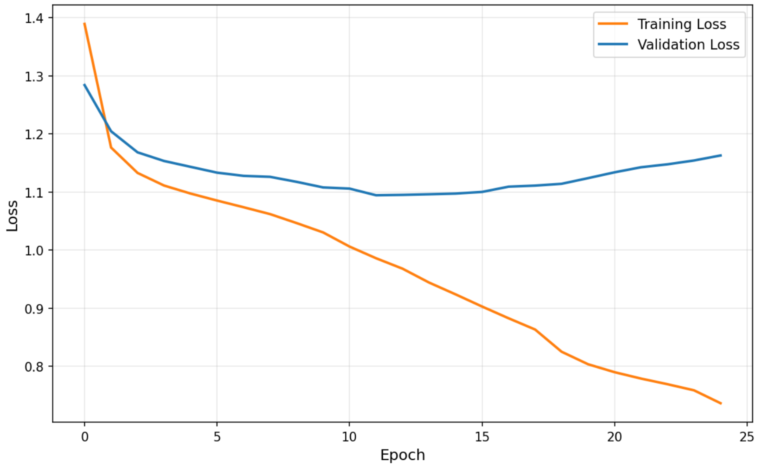


A

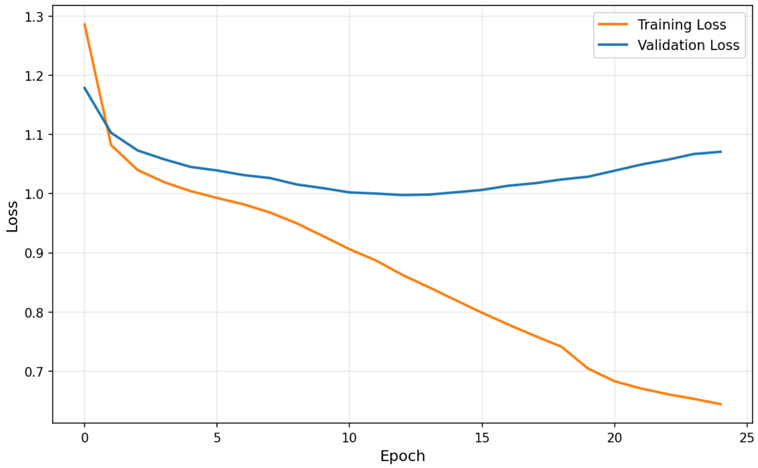


B

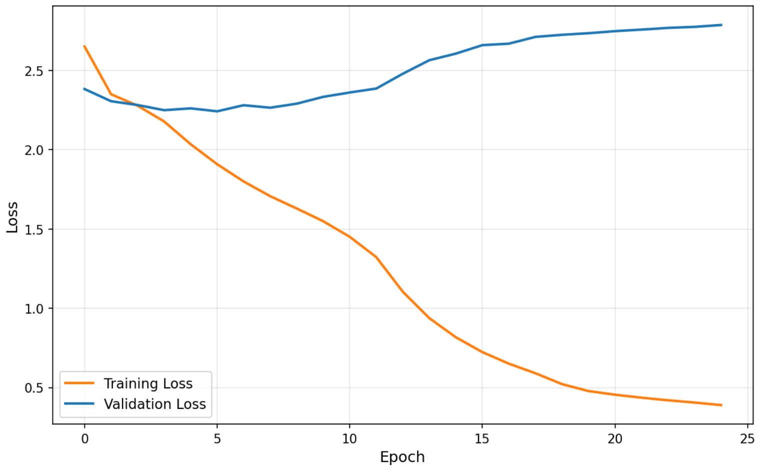


C

Figure S1: **Training loss curves for scWeave.** (A) Trained on the HiRES-embryo, HiRES-brain, CHARM-brain and GAGE-seq-brain datasets. (B) Trained on the HiRES-embryo, HiRES-brain, CHARM-brain, GAGE-seq-brain and four out of the six timepoints from the LiMCA dataset (C) Trained from scratch on the GAGE-seq-bone-marrow dataset.
